# Alternative cGAS signaling promotes herpes simplex encephalitis

**DOI:** 10.1101/2024.11.15.623778

**Authors:** Liraz Shmuel-Galia, Zhaozhao Jiang, Laurel Stine, Sara Cahill, Sze-Ling Ng, Ruth Wilson, Richard Kumaran Kandasamy, Evelyn A. Kurt-Jones, Joshi M. Ramanjulu, John Bertin, Viera Kasparcova, G. Scott Pesiridis, Katherine A. Fitzgerald, Fiachra Humphries

## Abstract

During infection, foreign DNA is sensed by cyclic GMP-AMP synthase (cGAS) leading to the production of cGAMP, STING-dependent type I interferon and proinflammatory cytokine expression, and autophagy. To prevent a response to self-DNA, cGAS activity is tightly regulated. Dysregulation of cGAS underpins interferonopathies, such as Aicardi-Goutières syndrome, as well as Lupus and neurodegenerative diseases like Parkinson’s disease. Thus, cGAS and its product cGAMP are therapeutic targets. However, if cGAS functions independently of cGAMP signaling is undefined. Here, we identified an alternative signaling pathway that cGAS engages independent of cGAMP synthesis. We demonstrate that alternative cGAS signaling promotes hyperexpression of CXCL1 and enhanced neutrophil recruitment that facilitates viral dissemination during herpes simplex encephalitis. Our study is the first report of an alternative cGAS response independent of cGAMP highlighting a previously uncharacterized scaffold function for cGAS.

## INTRODUCTION

Cyclic GMP-AMP synthase (cGAS) is a crucial component of the innate immune system that plays a vital role in detecting and responding to viral and bacterial infections, as well as other cellular stressors (*1*). cGAS localizes to both the cytoplasm and the nucleus of cells. Upon infection or cellular damage, foreign or aberrant self-DNA is released into the cytosol and binds to cGAS via its DNA-binding domain (*2*). Upon binding to dsDNA, cGAS undergoes a conformational change that allows it to catalyze the synthesis of cyclic GMP-AMP (cGAMP) from ATP and GTP (*3*). The enzymatic activity of cGAS results in the formation of a second messenger molecule called cGAMP, which consists of a cyclic dinucleotide with a unique 2’,3’-cGAMP linkage (*2*). Synthesized cGAMP can then bind to and activate the endoplasmic reticulum (ER)-associated protein stimulator of interferon genes (STING) (*4*). STING then undergoes a conformational change, leading to its activation, oligomerization, and translocation to the Golgi where it recruits TANK-binding kinase 1 (TBK1) through its cytoplasmic tail (*5*). TBK1 then phosphorylates both STING and itself, which further propagates the signal and leads to the recruitment of the transcription factor interferon (IFN) regulatory factor 3 (IRF3)(*6*). Following phosphorylation, IRF3 translocates into the nucleus and binds to specific DNA sequences known as IFN-stimulated response elements to induce production of type I IFNs, such as IFN-β. cGAS-STING activation also results in the activation of nuclear factor κB (NFκB) and autophagy responses (*7*).

Numerous mechanisms exist to prevent self-DNA recognition by cGAS and cGAMP synthesis (*8*). In addition, viruses have evolved inhibitory proteins to target cGAMP synthesis during infection to suppress antiviral immune responses (*9-11*). cGAS was thought to localize exclusively to the cytosol where compartmentalization of nuclear or mitochondrial DNA precluded access to cGAS. Additionally, TREX1 (a DNA exonuclease) limits the accumulation of cytosolic dsDNA as another mechanism to limit cGAS activity. cGAS is predominantly nuclear in many cell types where it associates with chromatin (*12, 13*). The barrier-to-autointegration factor 1 (BAF) binds to cGAS and outcompetes it for DNA binding, thus, limiting cGAS activation by nuclear DNA (*14*). Recent cryogenic electron microscopy studies have uncovered the mechanism that facilitates nucleosome-mediated inhibition of cGAMP synthesis (*14-19*). Indeed, when tightly bound to nucleosomes, cGAS is locked in a monomeric state, whereby steric hindrance prevents cGAS activation and dimerization by genomic DNA.

Several monogenic diseases in humans have been defined that result from self-DNA engagement of the cGAS-STING pathway. One of these diseases, Aicardi-Goutières syndrome (AGS) is an early-onset neurological disorder characterized by constitutive production of type I IFNs and IFN-stimulated genes (*20, 21*). The plasticity of innate immune signaling pathways results in alternative signaling events that compensate for the inactivation of key effector molecules. Given the recent evidence of mechanisms that control cGAS activity independent of cGAMP synthesis in the nucleus, we hypothesized that cGAS may signal via its DNA-binding domain independently of cGAMP.

The cGAS catalytic domain is a target for viral inhibition; indeed, several viruses have evolved methods for inhibiting cGAS catalytic activity limiting cGAMP synthesis(*9*). In addition, pharmacological strategies targeting the cGAS catalytic site block cGAS activity to limit conditions such as AGS and Lupus. In this study, we generated mice expressing catalytically dead cGAS, where Glutamine 211 and Aspartic acid 213 were mutated to alanine (cGAS^E211A/D213A^). cGAS^E211A/D213A^ mice were unable to generate cGAMP in response to DNA binding. cGAS^E211A/D213A^ mice failed to produce cGAMP and had impaired responses to herpes simplex virus 1 (HSV-1) and dsDNA. Interestingly, cGAS^E211A/D213A^ mice were hypersusceptible to HSV-1 infection when compared to cGAS^-/-^ mice. In response to HSV-1, cGAS^E211A/D213A^ mice hyperproduced CXCL1 and had elevated neutrophil recruitment in the central nervous system (CNS). Neutralization of CXCL1 protected cGAS^E211A/D213A^ mice from HSV-1-induced lethality.

Our study identifies a cGAMP-independent function for cGAS whereby the DNA-binding domain of cGAS promotes CXCL1 expression to facilitate neutrophil recruitment in the brain, wider tissue damage, and dissemination of neuroinvasive strains of HSV-1 through the CNS. Our study highlights the consequences of viral immune evasion and identifies important considerations for therapeutic targeting of the cGAS catalytic domain.

## RESULTS

### Generation and evaluation of catalytically inactive cGAS mice

To evaluate cGAMP-independent functions of cGAS, we generated catalytically inactive cGAS mice by mutating two residues essential for cGAMP synthesis, E211 and D213 to alanine (**Supplementary Fig. 1a**). Mutation of these sites completely abrogated cGAMP synthesis from bone marrow-derived macrophages (BMDMs) exposed to transfected dsDNA (**Supplementary Fig. 1b)** but had no effect on exogenously delivered cGAMP-driven STAT1 phosphorylation or cGAS protein levels (**Supplementary Fig. 1c**). We also evaluated responses to a dsDNA virus, the Baccman virus (an insect virus), which induced IFNβ, CXCL10, and IFNα protein production, all of which were completely impaired in serum from cGAS^E211A/D213A^ mice 6 hours post-infection (**Supplementary Fig. 1d–f**), indicating that the cGAS-cGAMP pathway was inactivated.

### cGAS^E211A/D213A^ mice are hypersusceptible to herpes simplex encephalitis

cGAS and STING are essential for limiting the neuroinvasive McKrae strain that causes herpes simplex encephalitis (HSE) (*22*). Indeed, deletion of cGAS or STING results in hyper-susceptibility to HSV-1 infection due to impaired activation of cGAS in microglia (*23, 24*). To evaluate HSV-1-induced responses, we infected wild-type (WT), cGAS^-/-^, and cGAS^E211A/D213A^ BMDMs with HSV-1 and assessed cytokine production. HSV-1 infection induced robust production of IFNβ, TNF-α, and the IFN-stimulated genes (ISGs) CXCL10 and CXCL9 in WT BMDMs, and this induction was completely abrogated in cGAS^-/-^ and cGAS^E211A/D213A^ BMDMs (**Fig. 1a–d**). As expected, infection with an RNA virus (Sendai virus) resulted in comparable cytokine production between WT, cGAS^-/-^, and cGAS^E211A/D213A^ cells (**Fig. 1a–d**). In line with these data, cGAS^-/-^ and cGAS^E211A/D213A^ cells also displayed impaired mRNA expression of *Ifnβ, Il6*, and the ISGs *Cxcl10, Ccl*5, *Cxcl9*, and *Isg15* in response to HSV-1 infection **(Fig. 1e–j)**. cGAS^-/-^ and cGAS^E211A/D213A^ BMDMs also displayed impaired HSV-1-induced IRF3, TBK1, and STAT1 phosphorylation when compared to WT BMDMs **(Fig. 1k)**.

**Figure 1:**
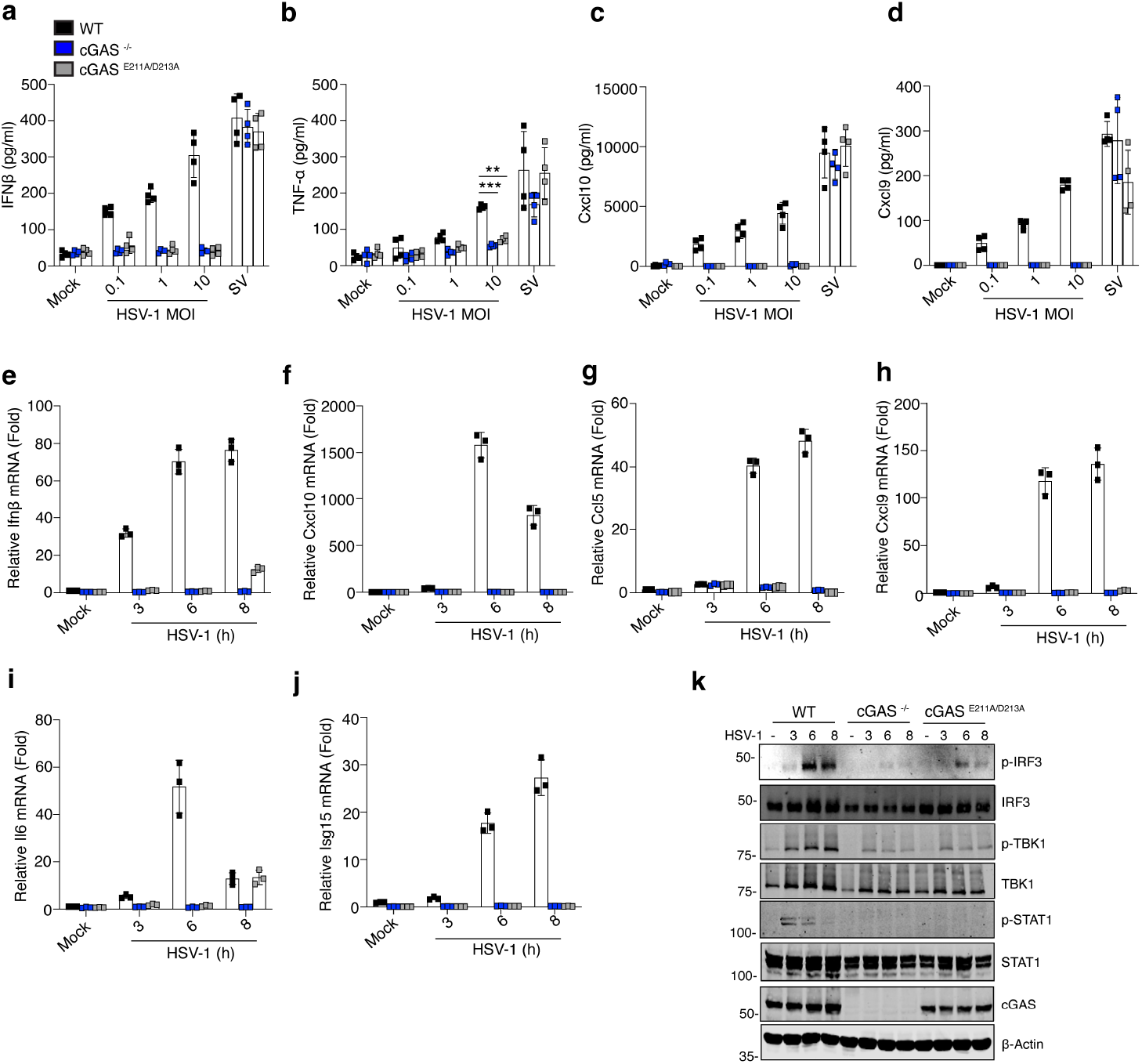
*Cgas*^E211A/D213A^ macrophages have impaired HSV-1 responses. (**a–d**) Bone marrow-derived macrophages (BMDMs) from wild-type (WT), *Cgas*^-/-^, and *Cgas*^E211A/D213A^ mice were mock infected with PBS (Mock) or infected with the indicated multiplicities of infection (MOI) of HSV-1 or with Sendai virus (SV, MOI=10) and the levels of the cytokines IFNβ (a), TNF-α (b), CXCL10 (c), and CXCL9 (d) were measured by ELISA. (**e–j**) Graphs showing the relative mRNA levels of *Ifnβ* (e), *Cxcl10* (f), *Ccl5* (g), *Cxcl9* (h), *Il-6* (i), and *Isg15* (j) compared to the housekeeping gene, TBP, as assessed by qRT-PCR in WT, *Cgas*^-/-^, and *Cgas*^E211A/D213A^ mice that were mock infected (Mock) or infected with HSV-1 (MOI =10) for the indicated timepoints. (**k**) Western blot showing the total protein levels and phosphorylated protein levels for IRF3, TBK1, and STAT1, as well as total cGAS in WT, *Cgas*^-/-^ and *Cgas*^E211A/D213A^ BMDMs infected with HSV-1 for the indicated time points (hours post-infection). β-Actin is shown as a loading control. The blot is representative of 3 experiments. A multiple comparison analysis was performed using a two-way ANOVA; ** *p*<0.01,*** *p*<0.001.

We next evaluated if cGAS^E211A/D213A^ mice displayed comparable sensitivities to cGAS^-/-^ mice when challenged with HSV-1 McKrae. WT, cGAS^-/-^, and cGAS^E211A/D213A^ mice were administered HSV-1 via ocular infection and monitored for clinical signs of HSE, including weight loss, hydrocephalus, survival, and neurological function. Interestingly, cGAS^E211A/D213A^ displayed increased weight loss **(Fig. 2a)**, hydrocephalus **(Fig. 2b)**, decreased survival **(Fig. 2c)** and elevated symptoms of neurological disease **(Fig. 2d)** following ocular infection. In addition, cGAS^E211A/D213A^ mice infected with HSV-1 had a significant increase in HSV-1 titer in the brain tissue 5 days post-infection **(Fig. 2e)**.

**Figure 2:**
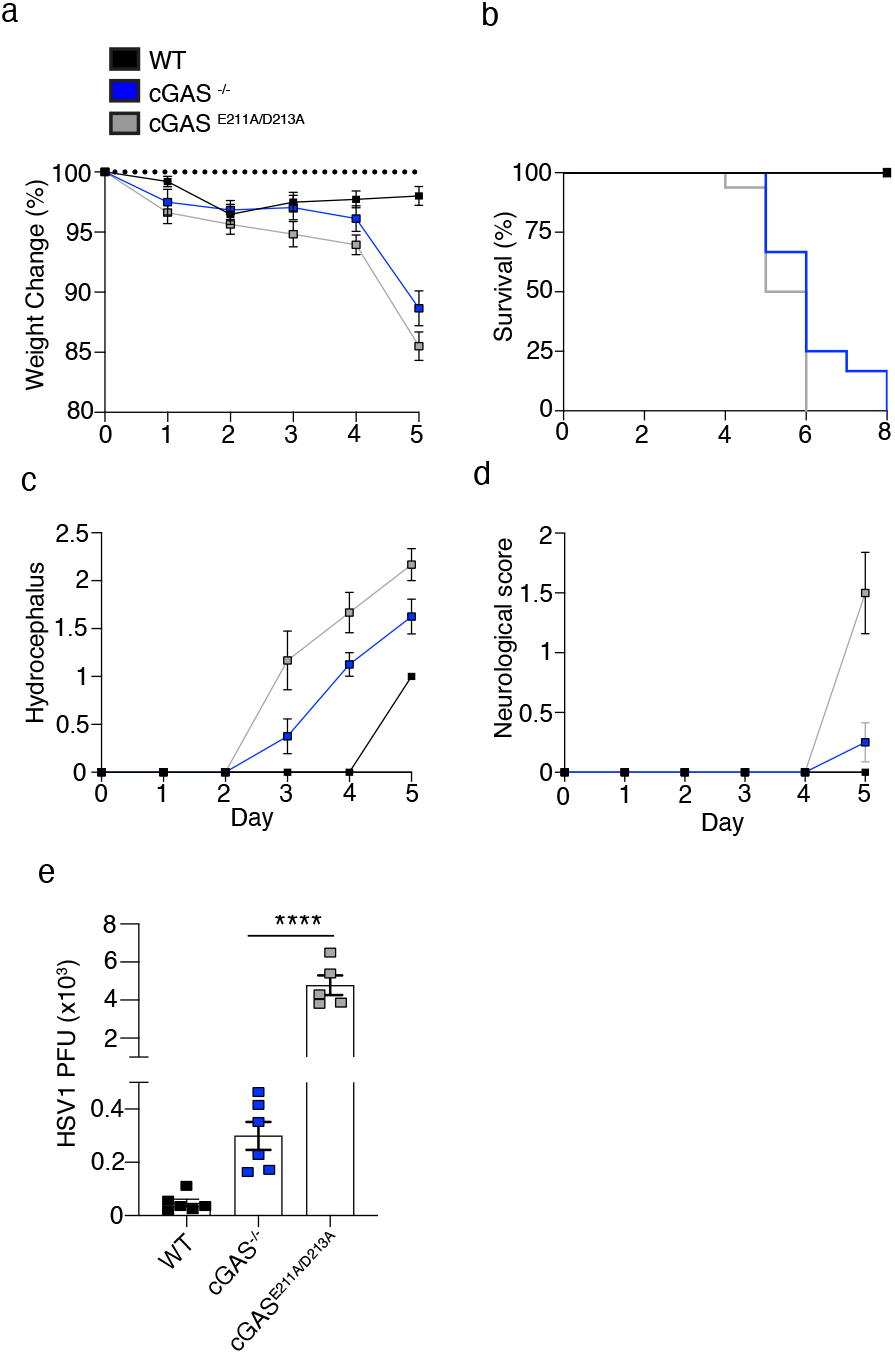
*Cgas*^E211A/D213A^ mice are hypersusceptible to neuroinvasive HSV-1 infection. (**a–d**) WT, *Cgas*^-/-^, and *Cgas*^E211A/D213A^ mice were mock infected with PBS (Mock) or infected with HSV-1 (2 × 10^5^ plaque-forming units [PFUs]) and monitored daily for weight change (a), survival (b), hydrocephalus (c), and neurological score (d) for up to 8 days (n =9-13 per group). The dotted line in A indicates baseline weight. (**e**) HSV-1 PFUs were measured in the brain tissue of wild-type, *Cgas*^-/-^, and *Cgas*^E211A/D213A^ mice infected with HSV-1 (2×10^5^ PFUs) at day 5 post-infection. For comparisons of two groups, a two-tailed Student’s t-test was performed. A multiple comparison analysis was performed using a two-way ANOVA; **** *p*<0.0001.

### cGAS^E211A/D213A^ induces hyperproduction of HSV-1-induced CXCL1

We next sought to define the molecular basis for the enhanced viral replication in cGAS^E211A/D213A^ mice when compared to cGAS^-/-^ mice. We hypothesized that catalytically inactive cGAS may play a structural role in eliciting an alternative signaling pathway in response to HSV-1 infection via its DNA-binding domain. Thus, we sequenced the transcriptome of dsDNA-induced WT cGAS^-/-^ and cGAS^E211A/D213A^ BMDMs to identify any transcriptional responses to DNA that were retained in the absence of cGAMP synthesis. Treatment of WT BMDMs with dsDNA led to a robust transcriptional response which resulted in the upregulation and downregulation of many antiviral and inflammatory genes. Both cGAS^-/-^ and cGAS^E211A/D213A^ BMDMs displayed a defective dsDNA-induced transcriptional response as expected **(Fig. 3a)**. Interestingly, cGAS^E211A/D213A^ cells displayed an increase in expression of Cxcl1 relative to cGAS^-/-^ cells. Cxcl1 was the only gene elevated in cGAS^E211A/D213A^ cells compared to cGAS^-/-^ cells **(Fig. 3a)**. We confirmed these findings by monitoring CXCL1 expression by qRT-PCR. dsDNA and HSV-1 infection induced CXCL1 expression in WT and cGAS^-/-^ cells. However, cGAS^E211A/D213A^ cells displayed a significant increase in dsDNA- or HSV-1-induced Cxcl1 expression compared to WT or cGAS^-/-^ cells **(Fig. 3b,c)**. Similarly, infection of cGAS^E211A/D213A^ BMDMs with HSV-1 resulted in a significant increase in CXCL1 protein secretion compared to HSV-1-infected WT or cGAS^-/-^ cells **(Fig. 3d)**. WT, cGAS^-/-^, and cGAS^E211A/D213A^ cells displayed comparable levels of Sendai virus-induced CXCL1 secretion **(Fig. 3d)**, demonstrating that the elevated response was specific to dsDNA stimulation.

**Figure 3:**
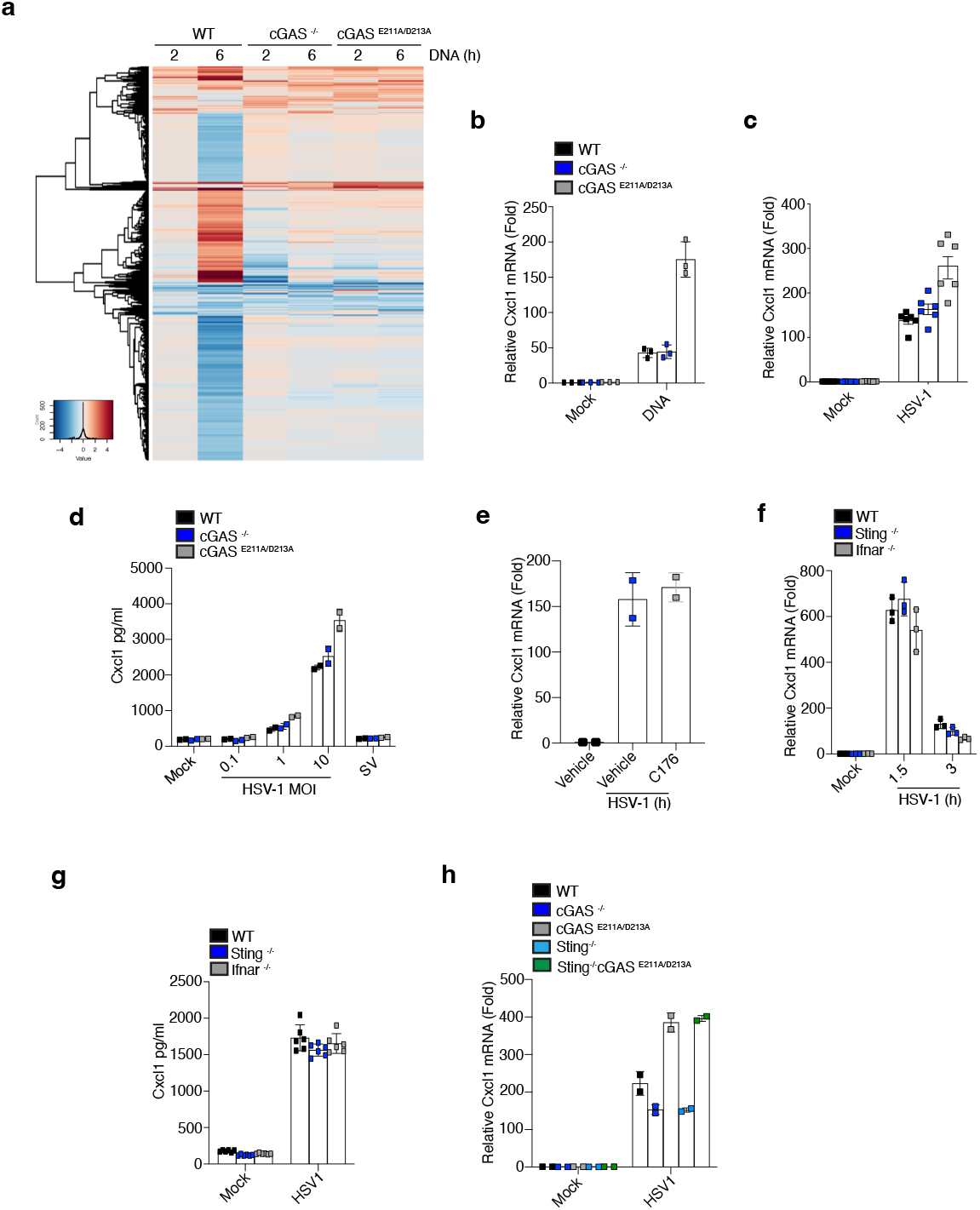
*Cgas*^E211A/D213A^ promotes hyperproduction of HSV-1-induced CXCL1 production. (**a**) Heatmap illustrating the differentially expressed genes in BMDMs from WT, *Cgas*^-/-^, and *Cgas*^E211A/D213A^ mice transfected with dsDNA (100ng) at the indicated time points. (**b**,**c**) WT, *Cgas*^-/-^, and *Cgas*^E211A/D213A^ BMDMs were uninfected, transfected with HSV DNA (b), or infected with HSV-1 (MOI =10) (c), and the level of Cxcl1 mRNA was assessed by qRT-PCR. (**d**) Protein levels of CXCL1 measured by ELISA in WT, *Cgas*^-/-^, and *Cgas*^E211A/D213A^ mice that were mock infected with PBS (Mock) or infected with the indicated MOIs of HSV-1 or SV (SV, MOI =1 for 24 hrs. (**e**) Cxcl1 mRNA expression assessed by qRT-PCR performed on BMDMs that were uninfected or infected with HSV-1 (MOI =10) and either untreated or treated with a STING inhibitor (C176). (**f**,**g**) BMDMs from WT, *Sting*^-/-^, and *Ifnar*^-/-^ mice were infected with HSV-1 (MOI =1) for the indicated time points and assessed for CXCL1 mRNA levels by qRT-PCR (**f**) and Cxcl1 protein levels by ELISA (g). (**h**) Cxcl1 mRNA levels assessed by qRT-PCR in WT, *Cgas*^-/-^, *Cgas*^E211A/D213A^, *Sting*^-/-^, and *Sting*^-/-^*Cgas*^E211A/D213A^ BMDMs. For comparisons of two groups, a two-tailed Student’s t-test was performed. A multiple comparison analysis was performed using a two-way ANOVA.

Given that cGAMP binds and activates STING, we hypothesized that STING may play a role in negatively regulating CXCL1 expression. However, inhibiting STING with C176, a small molecule STING inhibitor(*8*), did not affect CXCL1 expression in HSV-1-infected WT BMDMs **(Fig. 3e)**. Similarly, WT, *Sting*^-/-^, or *Ifnar*^-/-^ BMDMs displayed comparable levels of HSV-1-induced CXCL1 expression and secretion **(Fig. 3f,g)**. To further validate this, we intercrossed cGAS^E211A/D213A^ mice with *Sting*^-/-^ mice and evaluated HSV-1-induced CXCL1 expression. cGAS^E211A/D213A^ displayed elevated CXCL1 expression compared to WT, cGAS^-/-^ and *Sting*^-/-^ cells. However, cGAS^E211A/D213A^ STING-deficient cells retained augmented HSV-1-induced CXCL1 expression **(Fig. 3h)**. Thus, HSV-1-induced CXCL1 expression occurs independently of STING.

### HSV-1-induced CXCL1 expression depends on TLR2 and glycolysis

We next aimed to understand the signaling events that trigger and augment CXCL1 expression in the absence of cGAMP. HSV-1 is a DNA virus that can be recognized by cGAS. However, HSV-1 contains other pathogen-associated molecular patterns that are detected by other pattern-recognition receptors. HSV-1 can be sensed by Toll-like receptor 2 (TLR2, via HSV-1 glycoproteins) (*25*), TLR3 (via RNA produced during HSV-1 replication)(*26*), and TLR9 (via viral DNA)(*27*). We first assessed the requirement of TLR2 in mediating HSV-1-induced CXCL1 expression (*25*). WT and cGAS^E211A/D213A^ BMDMs were pre-treated with a TLR2 inhibitory peptide (TLR2p) (*28*). Inhibition of TLR2 impaired HSV-1-induced CXCL1 expression in both WT and cGAS^E211A/D213A^ BMDMs **(Fig. 4a)**. As the adaptor protein MyD88 is activated downstream of TLR2, we next evaluated the effect of MyD88 deficiency on HSV-1-induced CXCL1 expression. MyD88-deficient cells also failed to induce CXCL1 in response to HSV-1 infection **(Fig. 4b)**. Because IL-17-induced CXCL1 occurs in a TAK1/NFκB-dependent manner (*29*), we assessed if HSV-1-induced expression of CXCL1 depends on TAK1. Small molecule inhibition of TAK1 potently blocked HSV-1-induced CXCL1 expression and secretion in WT cells **(Fig. 4c,d**). In addition, TAK1 inhibition also impaired the augmented expression of CXCL1 in HSV-1-infected cGAS^E211A/D213A^ BMDMs **(Fig. 4e)**. Thus, HSV-1-induced expression of CXCL1 is dependent on a TLR2, MyD88, TAK1 signaling axis that occurs independently of cGAMP synthesis.

**Figure 4:**
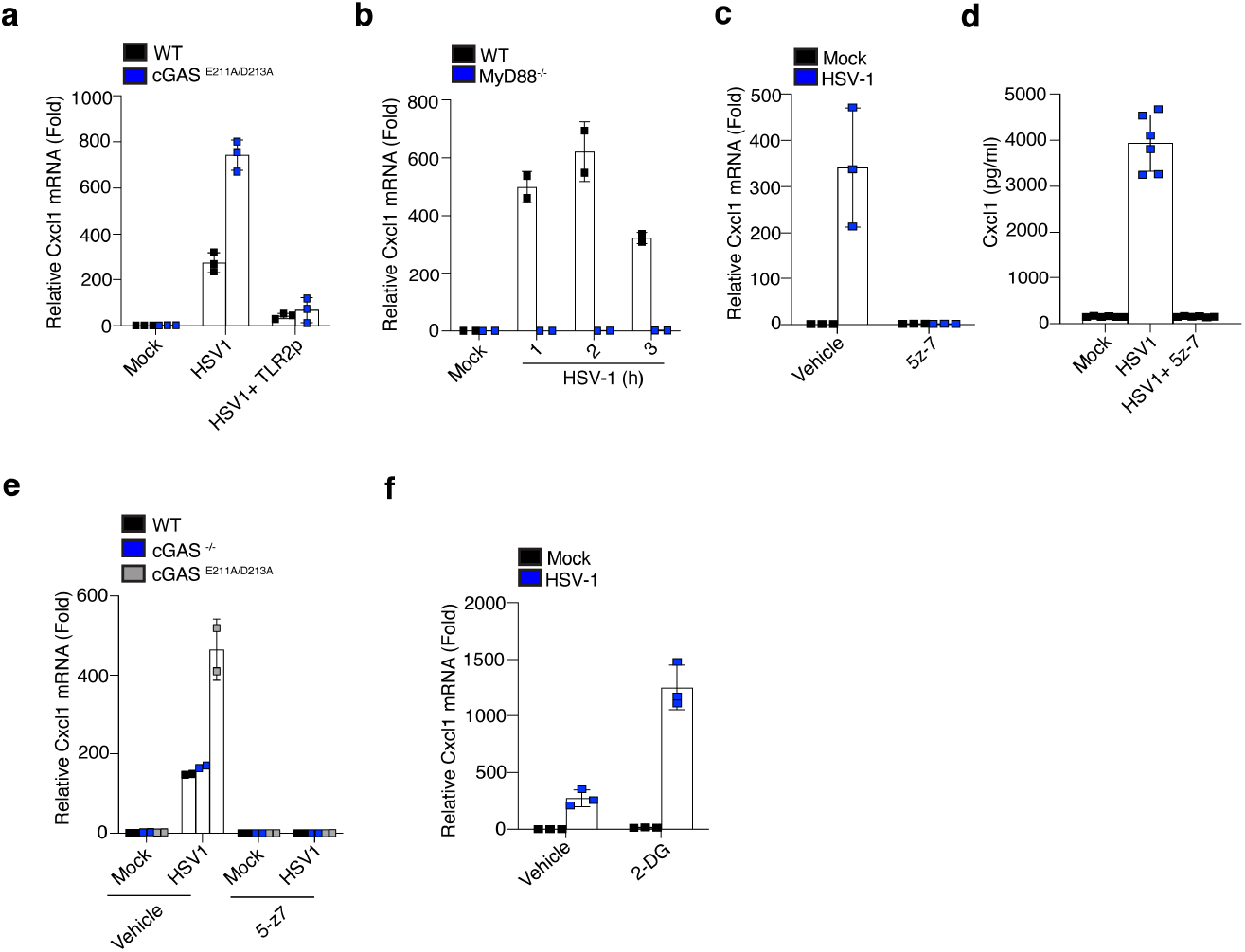
HSV-1-induced CXCL1 expression is dependent on TLR2 and OxPhos. (**a**) WT and *Cgas*^E211A/D213A^ BMDMs were pre-treated with a TLR2 inhibitory peptide (TLR2p) and assessed by qRT-PCR for Cxcl1 mRNA. (**b**) WT and MyD88^-/-^ BMDMs were mock infected with PBS (Mock) or infected with HSV-1 (MOI =10) and Cxcl1 mRNA was measured at the indicated time points by qRT-PCR. (**c**,**d**) WT BMDMs were mock infected with PBS (Mock) or infected with HSV-1 (MOI =10) and then treated with DMSO (Vehicle) or the TAK inhibitor (5z-7) and the Cxcl1 mRNA levels were assessed by qRT-PCR (c) and the CXCL1 protein levels were assessed by ELISA (d). (**e**) WT, *Cgas*^-/-^, and *Cgas*^E211A/D213A^ BMDMs were mock infected with PBS (Mock) or infected with HSV-1 (MOI =1) and untreated (Vehicle) or treated with the TAK inhibitor (5-z7) and Cxcl1 mRNA levels were assessed by qRT-PCR. (**f**) WT BMDMs were mock infected with PBS (Mock) or infected with HSV-1 (MOI=10) and untreated or treated with a glycolysis inhibitor (2-DG) and Cxcl1 mRNA levels were assessed by qRT-PCR. All qRT-PCR results are shown relative TBP. For comparisons of two groups, a two-tailed Student’s t-test was performed. A multiple comparison analysis was performed using a two-way ANOVA.

Given that HSV-1-induced CXCL1 expression depends on recognition of HSV-1 by TLR2, we hypothesized that the augmented CXCL1 expression observed in the cGAS^E211A/D213A^ cells occurred via a two-signal model whereby TLR ligation provides signal 1 and the absence of cGAMP synthesis and presence of the cGAS DNA-binding domain serves as signal 2 and engages an unknown pathway that is inhibited upon cGAMP synthesis. Because HSV-1 induces glycolysis in infected cells, we hypothesized that an alteration in cellular metabolism accounts for the elevation in CXCL1 transcription (*30*). To test our hypothesis, we used the glycolysis inhibitor 2-deoxyglucose (2-DG) to determine if inhibiting glycolysis regulates HSV-1-induced CXCL1 expression. Interestingly, pre-treatment of WT BMDMs with 2-DG resulted in a significant increase in HSV-1-induced CXCL1 expression when compared to vehicle-treated cells **(Fig. 4f)**. Thus, HSV-1-induced glycolysis negatively regulates CXCL1 expression.

### CXCL1 blockade protects against herpes simplex encephalitis

CXCL1 is a chemokine whose expression has been implicated in several inflammatory diseases. CXCL1 plays an important role in promoting angiogenesis and is essential for recruiting neutrophils (*31*). In addition, neutrophil recruitment to the CNS has been implicated as a key driver of HSE and viral dissemination due to tissue damage (*32*). Thus, we next assessed if cGAS^E211A/D213A^ mice displayed enhanced CXCL1 production and neutrophil recruitment during HSV-1 infection *in vivo*. Intraperitoneal infection of HSV-1 resulted in CXCL1 production in the serum and an increase in peritoneal neutrophils 6 hours post-infection in WT and cGAS^-/-^ mice. However, cGAS^E211A/D213A^ mice produced a significant increase in serum CXCL1 and recruitment of Ly6G^+^ positive neutrophils in response to HSV-1 infection when compared to WT and cGAS^-/-^ mice **(Fig. 5a,b)**. In addition, ocular infection of HSV-1 also significantly increased CXCL1 expression in the trigeminal ganglion tissue 24 hours post-infection **(Fig. 5c)**. Based on these data, we hypothesized that the elevated CXCL1 production induced by HSV-1 promotes more tissue damage and viral dissemination, which may account for the elevated viral replication we observed in the brain tissue of cGAS^E211A/D213A^ mice **(Fig. 2e)**. Thus, we used an anti-CXCL1 neutralizing antibody to evaluate if blocking CXCL1 alleviates the augmented HSE observed in the cGAS^E211A/D213A^ mice. Compared to an isotype control, anti-CXCL1 treatment had marginal effects on weight loss in cGAS^E211A/D213A^ mice **(Fig. 5d)**. However, anti-CXCL1-treated HSV-1-infected cGAS^E211A/D213A^ mice showed improved survival and a significant decrease in the development of hydrocephalus when compared to isotype control-treated cGAS^E211A/D213A^ mice **(Fig. 5e,f)**. Our results demonstrate that increased CXCL1 expression is involved in HSE pathogenesis.

**Figure 5:**
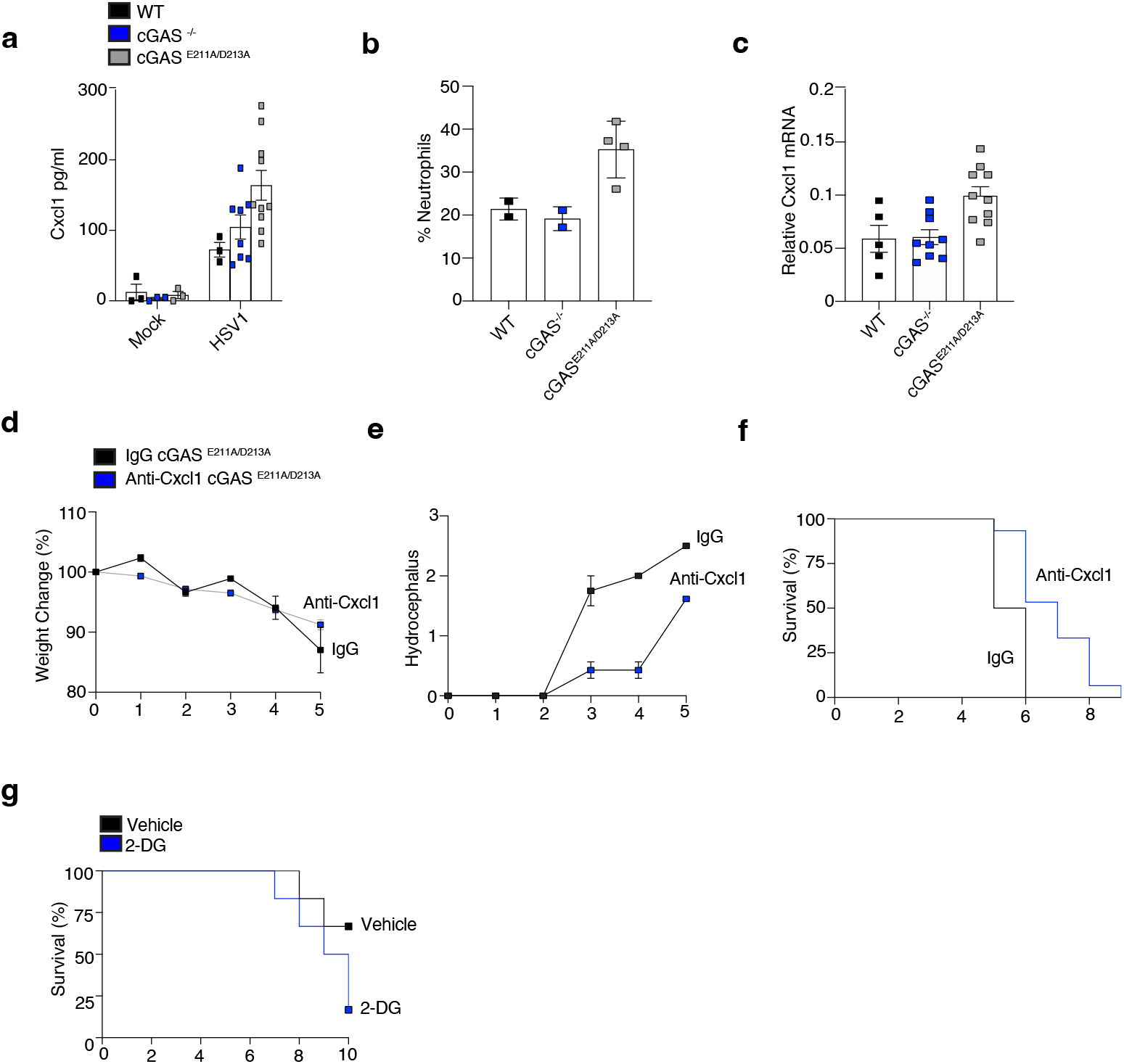
Modulation of CXCL1 levels regulates the severity of herpes simplex encephalitis. (**a**,**b**) WT, *Cgas*^-/-^, and *Cgas*^E211A/D213A^ mice were mock infected with PBS (Mock) or infected with HSV-1 intraperitoneally and Cxcl1 protein levels in the serum were measured by ELISA (a) and Ly6G+ neutrophils were assessed by FACS (b). (**c**)WT, *Cgas*^-/-^, and *Cgas*^E211A/D213A^ mice were infected with HSV-1 (2×10^5^ PFUs) via the ocular route and Cxcl1 mRNA levels were measured by qPCR. (**d–f**) *Cgas*^E211A/D213A^ mice were infected as in (c) and treated with an anti-Cxcl1 antibody or an isotype control (IgG) and weight change (d), hydrocephalus (e), and survival (f) measurements were taken at the indicated time points. (**g**) Survival rate in WT mice infected with the McKrae HSV-1 strain to induce HSE and treated with vehicle or a glycolysis inhibitor (2-DG).

We next sought to examine the effects of blocking HSV-1-induced glycolysis on HSE. Our in vitro findings demonstrated that 2-DG inhibition of the glycolytic shift induced by HSV-1 infection elevates CXCL1 expression. Thus, we evaluated the effects of 2-DG on HSE in the HSV-1 McKrae neuroinvasive ocular infection model. Infection of WT mice with HSV-1 McKrae was self-limiting in the majority of mice. However, WT mice administered daily doses of 2-DG were highly susceptible to HSE infection as evidenced by a significant reduction in the survival rate following infection **(Fig. 5g)**. Thus, blocking glycolysis increases HSE susceptibility in mice.

Altogether, these data demonstrate that, in the absence of cGAMP synthesis, catalytically inactive cGAS retains a scaffolding function via its DNA-binding domain that synergizes with TLR2 signaling to promote CXCL1 expression. During HSV infection, increased glycolysis produces CXCL1, and blocking glycolysis decreases survival in an HSE mouse model.

## DISCUSSION

Binding of cGAS to pathogen- or host-derived dsDNA promotes a conformational change activating cGAS catalytic activity and the conversion of ATP and GTP into the non-canonical cyclic dinucleotide cGAMP (*2, 3*). cGAMP then binds and activates STING to promote the production of antiviral responses (*3, 33*). However, aberrant activation of cGAS or STING can result in the onset of monogenic diseases known as interferonopathies (*21, 34, 35*). In addition, tissue damage associated with a number of autoinflammatory diseases can trigger the release of host DNA which stimulates cGAS activity and augments inflammation. Given the damaging consequences of excessive cGAMP synthesis, several regulatory mechanisms restrict cGAS binding to host DNA.

cGAS is localized to both the cytosol and nuclear compartment in close proximity to genomic DNA (*13*). In the nucleus, cGAS is tightly bound to chromatin and locked in an inactive state whereby steric hindrance prevents binding to genomic DNA (*13, 14, 16-18*). Indeed, certain forms of AGS exhibit disruption in the nuclear tethering of cGAS, altering its localization (*34*). Further, the cGAS catalytic site can be targeted by viral proteins to evade host responses. Given its importance, cGAMP synthesis now represents a viable therapeutic strategy to inhibit cGAS activity in disease (*20*).

Due to the plasticity of innate immune responses and compensatory mechanisms that exist within inflammatory pathways, we investigated if cGAS retains any functional role independent of cGAMP synthesis. To evaluate this, we generated a double-point mutant mouse in the catalytic domain of cGAS. Cells from these mice failed to produce cGAMP in response to DNA and displayed defective DNA- and viral-induced type I IFN responses. However, the protein levels of cGAS^E211A/D213A^ were comparable to WT cGAS. Interestingly, cGAS^E211A/D213A^ mice were hypersusceptible to a neuroinvasive strain of HSV-1 and developed more severe HSE compared to WT or cGAS^-/-^ mice. These findings indicated that catalytically inactive cGAS may engage an alternative signaling mechanism during HSV-1 infection. As HSV-1 is a DNA virus, it is likely that this mechanism occurs via the cGAS DNA-binding domain. Our future studies are now focused on understanding the direct contribution of the cGAS DNA-binding domain in this alternative signaling pathway.

Given that the enhanced CXCL1 expression by cGAS^E211A/D213A^ occurred independently of STING, we hypothesized that an alternative PRR was being triggered by HSV-1. Indeed, TLR2 was required for CXCL1 expression in response to HSV-1. This raised the possibility that cGAS^E211A/D213A^ synergizes with TLR2 signaling to promote CXCL1 expression in a transcriptional-independent manner. Glycolysis has been previously shown to limit HSV-1 replication and the development of HSE due to defective T-cell responses (*30*). However, it’s possible that enhanced *CXCL1* expression also contributes to HSE when glycolysis is blocked. Thus, we hypothesized that the recognition of HSV-1 DNA by cGAS may alter the metabolic state of the cell resulting in elevated expression of CXCL1. Indeed, 2-DG promoted HSV-1-induced CXCL1 expression and sensitized WT mice to HSV-1-induced HSE. Thus, we propose a two-signal model whereby initiation of the transcriptional response via TLR2 initiates CXCL1 expression that is augmented by the recognition of HSV-1 DNA by cGAS. Our future work is now focused on delineating the precise molecular details underpinning the alternative cGAS signaling during HSV-1 infection.

In summary, we have identified an alternative cGAS signaling pathway, triggered by DNA virus infection when cGAMP synthesis is impaired. When the catalytic activity of cGAS is blocked, HSV-1 infection elicits a rapid expression of CXCL1 resulting in enhanced neutrophil recruitment, tissue damage, and viral dissemination in the CNS.

**Supplementary Table 1.**
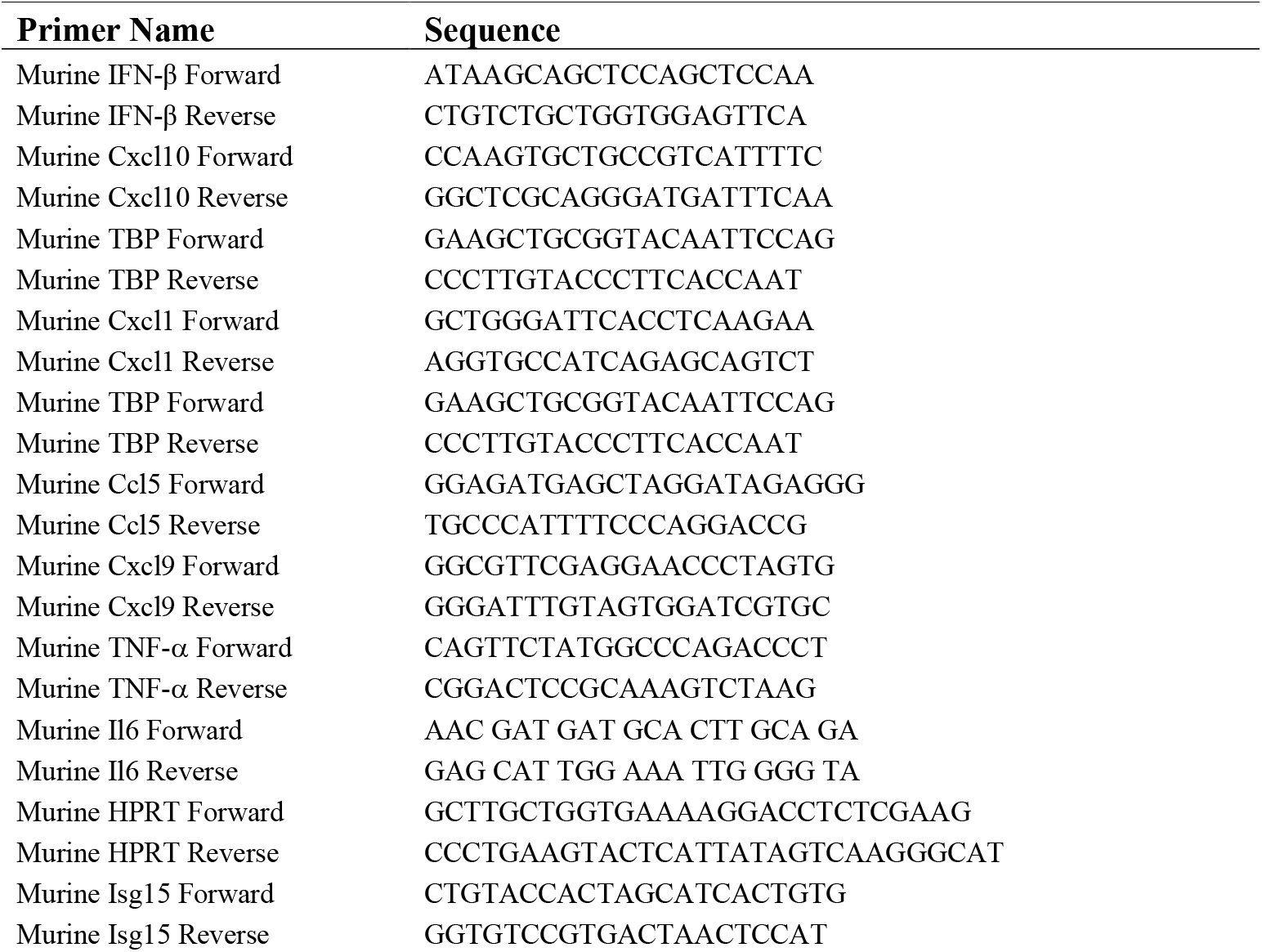
List of primer sequences used.

**Supplementary Figure 1:**
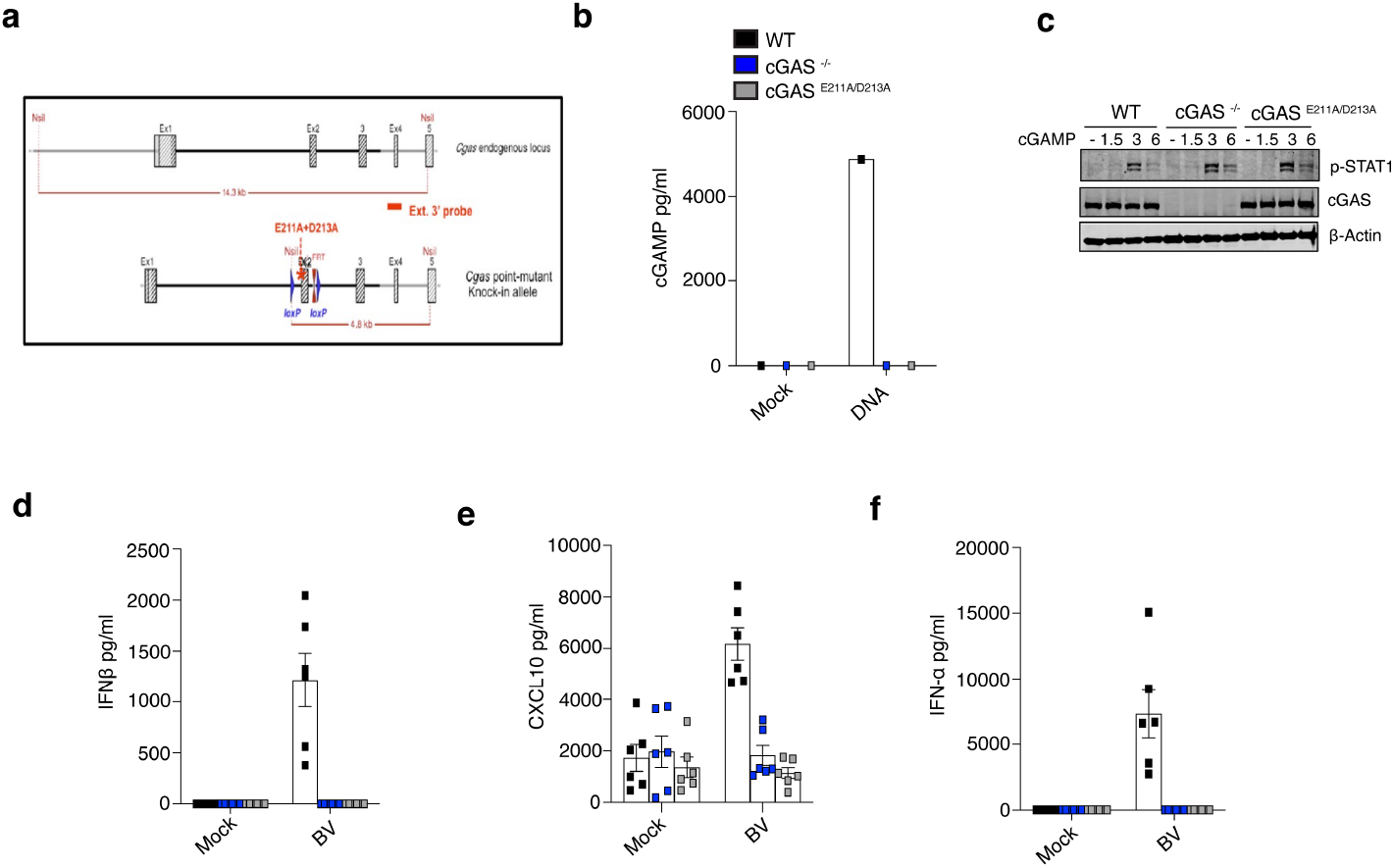
Generation and characterization of *Cgas*^E211A/D213A^ mice. (**a**) Schematic of the strategy used to generate the *Cgas*^E211A/D213A^ mice. (**b**) cGAMP levels assessed by ELISA in BMDMs from WT, *Cgas*^-/-^, and *Cgas*^E211A/D213A^ mice transfected with dsDNA (100ng). (**c**) Western blot to detect phosphorylated STAT1 and cGAS in WT, *Cgas*^-/-^, and *Cgas*^E211A/D213A^ mice treated with the indicated timepoints. (**d–f**) WT, *Cgas*^-/-^, and *Cgas*^E211A/D213A^ mice infected intraperitoneally with BK virus and the serum levels of IFNβ (d), Cxcl10 (e), and IFN-α (f) were measured by ELISA at 6 hours post-infection (n = 6 for each group). A multiple comparison analysis was performed using a two-way ANOVA.

## MATERIALS AND METHODS

### Biosafety

All study protocols were reviewed and approved by Environmental Health and Safety at UMass Chan Medical School prior to study initiation. All experiments with HSV-1 McKrae were performed in a biosafety level 2 laboratory by personnel equipped with the appropriate PPE.

### Mice

Animals were kept in a specific pathogen-free environment. *Cgas*^-/-^, *Sting*^-/-^, and *Ifnar*^-/-^ mice were used as previously described (*6*). *Cgas*^E211A/D213A^ mice were generated by GenOway using Flp recombination. *Cgas*^E211A/D213A^ was expressed under the control of the endogenous cGAS promoter. *Cgas*^+/E211A/D213A^ heterozygous mice were intercrossed to generate *Cgas*^E211A/D213A^ homozygous mice. *Sting*^-/-^ *Cgas*^E211A/D213A^ mice were generated by crossing *Sting*^-/-^mice with *Cgas*^E211A/D213A^ mice. Animals were housed in groups and fed standard chow diets. Sample sizes used are in line with other similar published studies.

### Culture of HSV-1 McKrae

T175 flasks of VeroE6 cells were infected with HSV-1 McKrae at an MOI of 0.1 for 48 hours. Supernatants were centrifuged at 450*g* for 10 minutes and aliquoted and stored at -80 ºC. Virus titre was determined by plaque assay in VeroE6 cells.

### HSV-1 McKrae infection of mice

The HSV-1 McKrae infections were performed as previously described (*6*). In brief, 8–12-week-old male and female mice were anesthetized with intraperitoneal injection of ketamine (100 mg kg^−1^ body weight) and xylazine (10 mg kg^−1^ body weight). Corneas were scratched in a 10 × 10 crosshatch pattern and mice were either inoculated with 2 × 10^5^ PFU of HSV-1 in 5 μl of PBS or mock infected with 5 μl of PBS. Mice were monitored daily for weight loss and assessed for ocular hair loss, eye swelling, hydrocephalus, and symptoms related to neurological disease as previously described (*23*).

### HSV-1 intraperitoneal infection of mice

Mice were infected intraperitoneally with HSV1 1×10^6^ PFU/mouse. Peritoneal cells were collected at the indicated times via peritoneal lavage using PBS.

### HSV-1 plaque assay

VERO cells were plated in 6 well plates (5×10^5^/well) in DMEM containing 10% FCS and 1% penicillin-streptomycin. The following day, brains were extracted from mice, homogenized in 1 ml of medium, and centrifuged for 10 min at 8000 *g*. The supernatant was collected and serially diluted 2-fold in DMEM. 500 μl of each dilution was added to the VERO cells for 1 h with gentle shaking. After 1 h, the medium was removed and replaced with 2 ml of DMEM containing 15 μg/ml IGS from human serum antibody (Sigma 14506). Cells were incubated at 37 °C for 2 days. Medium was then removed, and cells were fixed in 100% ice-cold methanol for 3 min. Plaques were stained with 10% crystal violet for 20 min with gentle shaking, washed, and counted.

### cDNA synthesis and real-time quantitative PCR

Total RNA was extracted from whole lung tissue or cells. 1 μg of RNA was reverse transcribed using the iScript cDNA synthesis kit (BioRad). 5 ng of cDNA was then subjected to qPCR analysis using iQ SYBR Green super-mix reagent (BioRad). Gene expression levels were normalized to a housekeeping gene (TATA-binding protein or HPRT). Relative mRNA expression was calculated by a change in cycling threshold method as 2^-ddC(t)^. The specificity of RT-qPCR amplification was assessed by a melting curve analysis. The sequences of primers used in this study are listed in **Supplementary Table 1**.

### Cell culture

For isolation of BMDMs, tibias and femurs were removed from WT mice, and bone marrow was flushed with complete DMEM-medium. Cells were plated in medium containing 20% (v/v) conditioned medium of L929 mouse fibroblasts cultured for 7 days at 37 °C in a humidified atmosphere of 5% CO_2_. Medium was replaced every 3 days.

### cGAMP assay

cGAMP was quantified by ELISA assay as per manufacturer’s instructions (Cayman Chemical). Briefly, plates were pre-washed 5 times with 300ul of wash buffer. 2’3’ cGAMP ELISA standard was serially diluted 8 times. 50ul of cell lysate from WT, *Cgas*^-/-^ and *Cgas*^E211A/D213A^ were added and incubated with the 2’3’-cGAMP-HRP tracer. Followed by incubation with the 2’3’ cGAMP polyclonal antiserum. The plate was then incubated overnight at4 °C. Plate was then washed 5 times with 300ul of wash buffer. 175ul of TMB substrate was added to each well for 30 minutes. Followed by addition of 75ul of stop solution. The plate was then read at a wavelength of 450nM. cGAMP levels were extrapolated from the standard curve.

### ELISA

Conditioned media or serum was collected as indicated and mouse IFNβ, TNF-α, CXCL10, CXCL9, and CXCL1 were quantified by sandwich ELISA (R&D Systems).

### Immunoblotting

For cell lysate analysis, cells were lysed directly in lysis buffer (50 mM Tris-HCl [pH 7.4], containing 150 mM NaCl, 0.5% [w/v] IgePal, 50 mM NaF, 1 mM Na_3_VO_4_, 1 mM dithiothreitol, 1 mM phenylmethylsulfonyl fluoride and protease inhibitor cocktail). Samples were resolved by SDS-PAGE, transferred to nitrocellulose membranes, and analyzed by immunoblot. Immunoreactivity was visualized by the Odyssey Imaging System (LICOR Biosciences). Anti-p-IRF3, anti-IRF3, anti-p-TBK1, anti-TBK1 anti-cGAS, anti-P-STAT1, anti-STAT1 antibodies were purchased from Cell Signaling Technology, Inc. Anti-β-actin (AC-15; A1978) was purchased from Sigma; anti-mouse IRDye™ 680 (926-68070) and anti-rabbit IRDye™ 800 (926-32211) were from LI-COR Biosciences.

### Flow cytometry

Isolated peritoneal cells were stained with anti-CD45.2 BV650, anti-CD11b BV510, anti-Ly6C APC, anti-Ly6G PE-cy7, anti-Ly6G FITC. Cells were acquired on a Cytek™ Aurora technology. Flow cytometry analysis was done with the FlowJo software.

### RNA sequencing

RNA sequencing was performed by BGI.

### Ethics

All animal studies were performed in compliance with the federal regulations set forth in the Animal Welfare Act, following the recommendations in the Guide for the Care and Use of Laboratory Animals of the National Institutes of Health, and the guidelines of the UMass Chan Medical School Institutional Animal Use and Care Committee (IACUC). All protocols used in this study were approved by the UMass Chan Medical School IACUC (protocol no. A-1633).

### Statistical analysis

For comparisons of two groups, a two-tailed Student’s t-test was performed. Multiple comparison analysis was performed using two-way ANOVA. 3 to 16 mice were used per experiment, sufficient to calculate statistical significance, and in line with similar studies published in the literature.

## Acknowledgments

We thank David Leib for providing the HSV-1 McKrae. We thank Melanie Trombly for assisting with manuscript preparation and editing. FH is supported by a Child Health Award from the Charles H. Hood Foundation. LSG is supported by the Crohn’s and Colitis Career Development Award.

## Author contributions

FH and LSG conceived the study, developed the concept, performed experiments, analyzed data, and wrote the manuscript. LS, RW, ZJ, and SN performed experiments and analyzed data. RKK analyzed RNA seq data and generated heat maps. EKJ provided advice on HSV-1 infections. JMR, VK, and JB provided the *Cgas*^E211A/D213A^ mice. KAF and GSP conceived the study, supervised the research, and edited the manuscript.

## Data and materials availability

All data in this paper are presented in the main text and supplementary text.

